# Nuclear speckle fusion via long-range directional motion regulates the number and size of speckles

**DOI:** 10.1101/347955

**Authors:** Jiah Kim, Kyu Young Han, Nimish Khanna, Taekjip Ha, Andrew S. Belmont

## Abstract

Although the formation of RNA-protein bodies has been studied intensively, their mobility and how their number and size are regulated are still poorly understood. Here, we show significant increased mobility of nuclear speckles after transcriptional inhibition, including long-range directed motion of one speckle towards another speckle, terminated by speckle fusion, over distances up to 4 um and with velocities between 0.2-1.5 μm/min. Frequently, 3 or even 4 speckles follow very similar paths, with new speckles appearing along the path followed by a preceding speckle. Speckle movements and fusion events contribute to fewer but larger speckles after transcriptional inhibition. These speckle movements are not actin-dependent, but occur within chromatin-depleted channels enriched with small granules containing the speckle-marker protein SON. Our observations suggest a mechanism for long-range, directed nuclear speckle movements, contributing to overall regulation of nuclear speckle number and size as well as overall nuclear organization.

## Introduction

Intracellular compartmentalization in eukaryotic cells is thought to represent a key evolutionary advance by which complex biochemical processes can be optimized and regulated within highly crowded cellular environments^1, 2, 3^. Compartmentalization is achieved not only with membrane-bound organelles but also through membrane-less “bodies”, including P-bodies and stress granules in the cytoplasm and nucleoli, nuclear speckles, Cajal, and PML bodies in the nucleus. Most membrane-less bodies are RNA-protein-rich complexes which function in transcription, RNA processing, and/or protein modification^4, 5^. Recent studies have led to the concept of RNA-protein bodies forming as the result of a phase transition between constituent RNA and proteins and surrounding cytoplasm or nucleoplasm through a demixing driven by interactions between low complexity sequences and multivalent molecules^6, 7, 8, 9^. RNA-protein bodies have been proven to have general physical characteristics of liquid droplets ^4, 5^. For example, P-granules undergo rapid assembly/disassembly, and exhibit dripping, fusion, and wetting by shearing force^10^, and nucleoli exhibit viscous fluid dynamics such as liquid bridge formation when they are ruptured^11^. Fluorescence recovery after photobleaching (FRAP) analysis of molecular dynamics showed that RNA-protein bodies continuously turnover many of their constituent molecules over a time-scale from a few seconds to a few minutes^12, 13, 14^. These microscopy observations suggest that these bodies are in a liquid or gel state with a constant flux of constituent molecules.

Despite a number of studies examining liquid properties and related behaviors of other cellular bodies, both the physical properties and dynamics of nuclear speckle bodies have not been well studied. Defined originally as interchromatin granule clusters (IGCs) by electron microscopy^15, 16^, nuclear speckles contain large clusters of ~20 nm RNP granules and are enriched in RNA processing factors and polyadenylated RNAs^17, 18, 19^. Nuclei typically contain 20-40 irregular shaped nuclear speckles varying in size from ~0.5 μm to several μm^20^.

Because “pure” liquid droplet bodies theoretically are predicted to merge into progressively fewer and larger droplets, a major question regarding cell bodies in general is how their number and size are regulated in the cell. Interestingly, with transcriptional inhibition nuclear speckles become rounder and larger with a reduction in their overall number^21, 22^. Live-cell microscopy revealed that normally speckles were relatively stationary, but displayed extension and dissociating particles. However, no peripheral dynamics were observed in the absence of transcription^23^. These previous studies did not address directly whether nuclear speckles also exhibit the general physical properties of RNA-protein bodies or how speckle morphology changes dynamically in terms of speckle numbers, size and shape.

Using the change in nuclear speckle morphology induced by transcriptional inhibition as a model system, here we studied the mobility and liquid-like behaviors of nuclear speckles. Consistent with previous observations, we observed a general change in morphology after transcriptional inhibition, with speckles becoming bigger and rounder. This was accompanied by an increase in speckle mobility within the nucleus.

Surprisingly this increase in speckle mobility was not random. Instead we observed repeated long-range, directional movement of multiple speckles over similar nuclear trajectories or “tracks” which extended over micrometer distances. These movements ended with fusion of the smaller, mobile speckles with larger, typically stationary speckles. These repeated fusion events contributed to the overall decrease in nuclear speckle number and increased mean speckle size as a function of time after transcriptional inhibition.

Perturbing normal actin polymerization did not prevent speckle long-range motion, the directionality of this motion, or speckle fusion events. Therefore, we excluded the possibility that actin filaments might provide “tracks” for this speckle movement. Combining live-cell imaging with correlative super-resolution light microscopy revealed speckles move within chromatin-depleted channels. Moreover, paths followed by more than one speckle as visualized by live-cell microscopy contained a concentration of small granules containing the speckle-marker SON within these DNA-depleted channels.

Our results are the first demonstration of repeated, long-range directed motion with nucleation and fusion of nuclear speckles in a live cell, revealing a previously unsuspected trafficking mechanism of nuclear speckles likely controlling speckle number and possibly nuclear localization. Overall, our results suggest that the distribution of nuclear speckles is significantly affected by translocation as well as dynamic nucleation and fusion of nuclear speckles, and possibly by changes in nuclear chromatin structure that may remove barriers to long-range speckle movements.

## Results

### Inhibition of RNA polymerase II transcription changes speckle morphology and increases overall speckle mobility

We used SON protein as a speckle marker to visualize nuclear speckles with high contrast. The SON protein showed the highest concentration in nuclear speckles versus nucleoplasm of any speckle markers we tested, with less distributed diffusely or in foci outside of nuclear speckles than other RS-domain proteins such as SC35 or ASF/SF1^19, 24^. The SON protein contains an RS domain and also 6 other tandem repeat regions unique to SON^25^, which may function as a scaffold for protein assembly^25^. The RS-domain is found in a large number of mRNA processing proteins, many enriched in nuclear speckles, as well as some proteins related to RNA pol 2 transcription^26^. Knockdown of SON results in a significant change in nuclear speckle morphology as visualized by immunostaining of SC35, another RS-domain containing speckle-marker protein^25^.

Attempts to express SON-EGFP from a cDNA plasmid transgene resulted in variegated and unstable transgene expression. To obtain stable expression, we instead transfected Chinese Hamster Ovary K1 (CHOK1) cells with a BAC transgene containing the full-length SON human genomic sequence engineered by BAC recombineering to express a SON-EGFP fusion protein^24^. Use of this ~200 kb BAC containing large 5’ and 3’ human genomic sequences flanking the SON gene enabled isolation of stable clones expressing uniform levels of EGFP-SON.

To study dynamics of speckles after transcriptional inhibition, we used 5,6-dichloro-1-β-ribofuranosyl benzimidazole (DRB). DRB blocks transcription via inhibition of kinases involved in transcription elongation^27, 28^. We confirmed in our CHO cell system previous work^21, 22^ that large speckles become brighter, rounder, and fewer in number after DRB addition (Fig. 1a, b).

**Figure 1.**
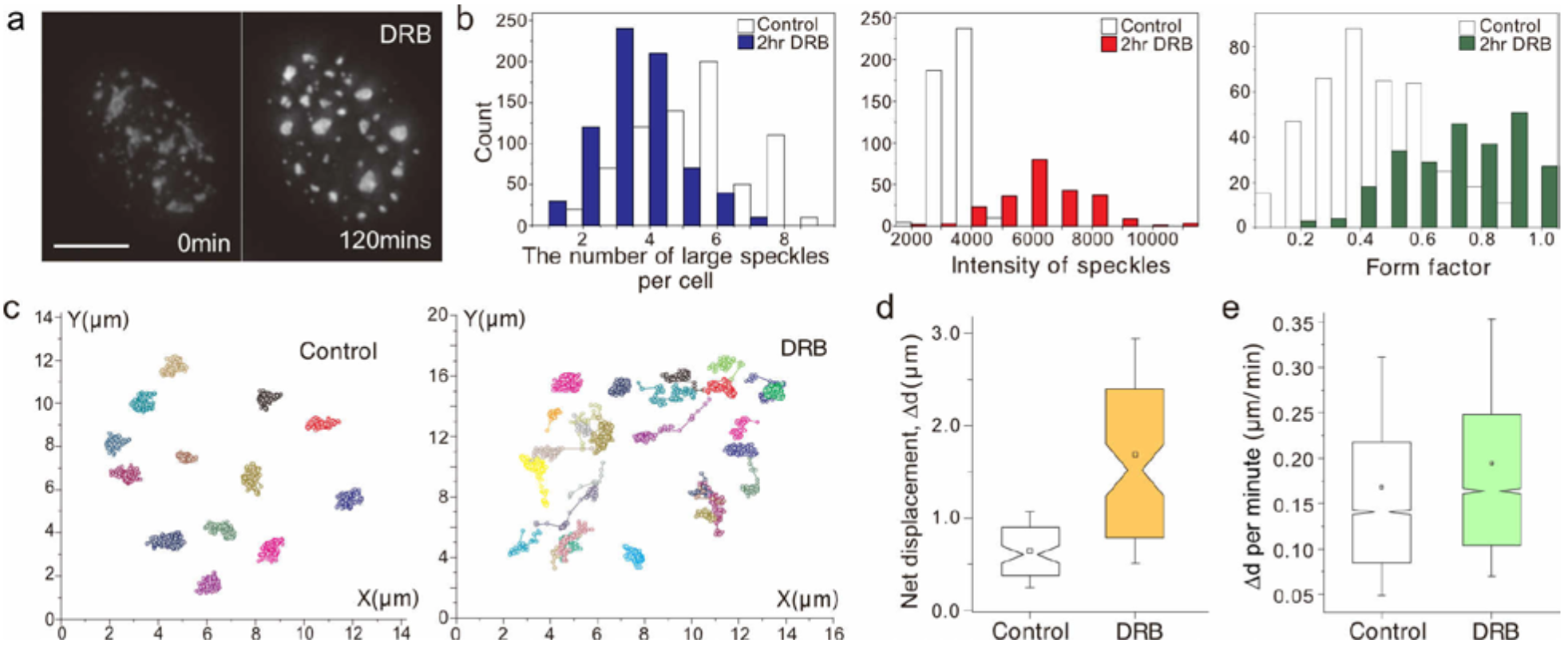
Inhibition of RNA polymerase II transcription by DRB changes speckle morphology and increases speckle mobility. **a.** Changes in speckle morphology, visualized using GFP-SON, before and after DRB addition. Scale bar: 5μm. **b.** Measurement of speckle number (>1 um in diameter) (left), intensity (middle), and shape (right) (Form Factor (FF)) before and after 2 hrs DRB treatment. Large speckles reduce in number (blue), become brighter (^~^2.5 fold, red), and rounder (1.5-fold increase in FF, green) after DRB addition. **c.** Speckle trajectories before and after DRB addition. Speckles, each assigned a different color, were tracked over a 1 hr period in control (left) and DRB-treated cell (right). **d.** Net displacement (Δd) of speckles measured in control (white) and DRB-treated cells (yellow). Speckle displacements increase after DRB addition (p-value = 1.007e-12; Paired student’s t-test). Boxplot: Box-mean (square inside box), median (notch of box), 25 (bottom) and 75 (top) percentiles; ends of error bars-10 (bottom) and 90 (top) percentiles. n=75 speckles from 10 control cells, n=80 speckles from 10 DRB treated cells. **e.** Displacement (Δd) per minute increases from before (white) to after DRB addition (green) (p-value=2.2e-16; paired Wilcoxon signed rank test) Boxplots as in (d). n=4487 steps from 49 speckles (control), n=4375 steps from 50 speckles (DRB).

The GFP-SON intensity within speckles became ~2.5-fold higher on average 2 hrs after DRB addition. The form factor (FF), defined as (4*π*Area/(Perimeter)^2^), increased ~50% from 0.50 in control cells to 0.76 on average in DRB treated cells. The FF ranges from 1 for a perfect circle to 0 for a line. Transcriptional inhibitors α-amanitin and triptolide (TRL) and the heavy metal cadmium(Cd) had similar effects on speckle morphology as DRB (Supplementary Fig. 1). However, we chose DRB for our imaging studies because it produced less rounding and change in shape of the nucleus in comparison to these other inhibitors. This reduced change in nuclear shape was critical for our tracking of speckle movement using alignment of nuclear images from different time points.

Live-cell imaging revealed an obvious increase in nuclear speckle mobility during DRB treatment. (Fig. 1c-e) To facilitate analysis, at each 1minute time point we used 2D projections of 3D z-stacks. We used a cross-correlation approach to best correct for nuclear rotations and translations between adjacent time points. In comparison to control cells, in which nuclear speckles showed restricted movements over 1 hour (Fig. 1c, left panel), in DRB-treated cells a significant fraction of nuclear speckles showed longer-range net displacements (Fig. 1c, right panel). Overlapping trajectories corresponded to examples in which nuclear speckles came together and merged (Fig. 1c). DRB treatment increased the mean net displacement, Δd, of nuclear speckles over 1.5 hour from 0.64 to 1.69 μm (p-value = 1.007e-12, paired t-test, Fig. 1d), with the increase of the mean distance change per 1 min time point, from 0.17 to 0.19 μm (p-value=2.2e-16, paired Wilcoxon signed rank test, Fig. 1e).

### DRB treatment induces long-range, directional speckle movements

We characterized movement of individual nuclear speckles after DRB addition by measuring the asymmetry coefficient, AC, together with the net displacement, Δd. We observed speckles for 1.5 hours beginning 30 mins after DRB addition using a 1 min time interval, tracking only those speckles which could be observed in at least 20 consecutive time points. To calculate the AC, we first measured the angle between the directions of speckle displacements for two consecutive time points (Fig. 2a). The AC was then defined as the logarithm to the base 2 of the ratio between the frequencies of backward (180 +/− 30^0^) versus forward (+/−30^0^) movements over the entire speckle trajectory^29^. Random diffusion produces AC values near 0, since forward motion and backward motion are equally likely. AC values higher than 0 indicate a bias towards forward movements, while AC values below 0 suggest constrained diffusion.

**Figure 2.**
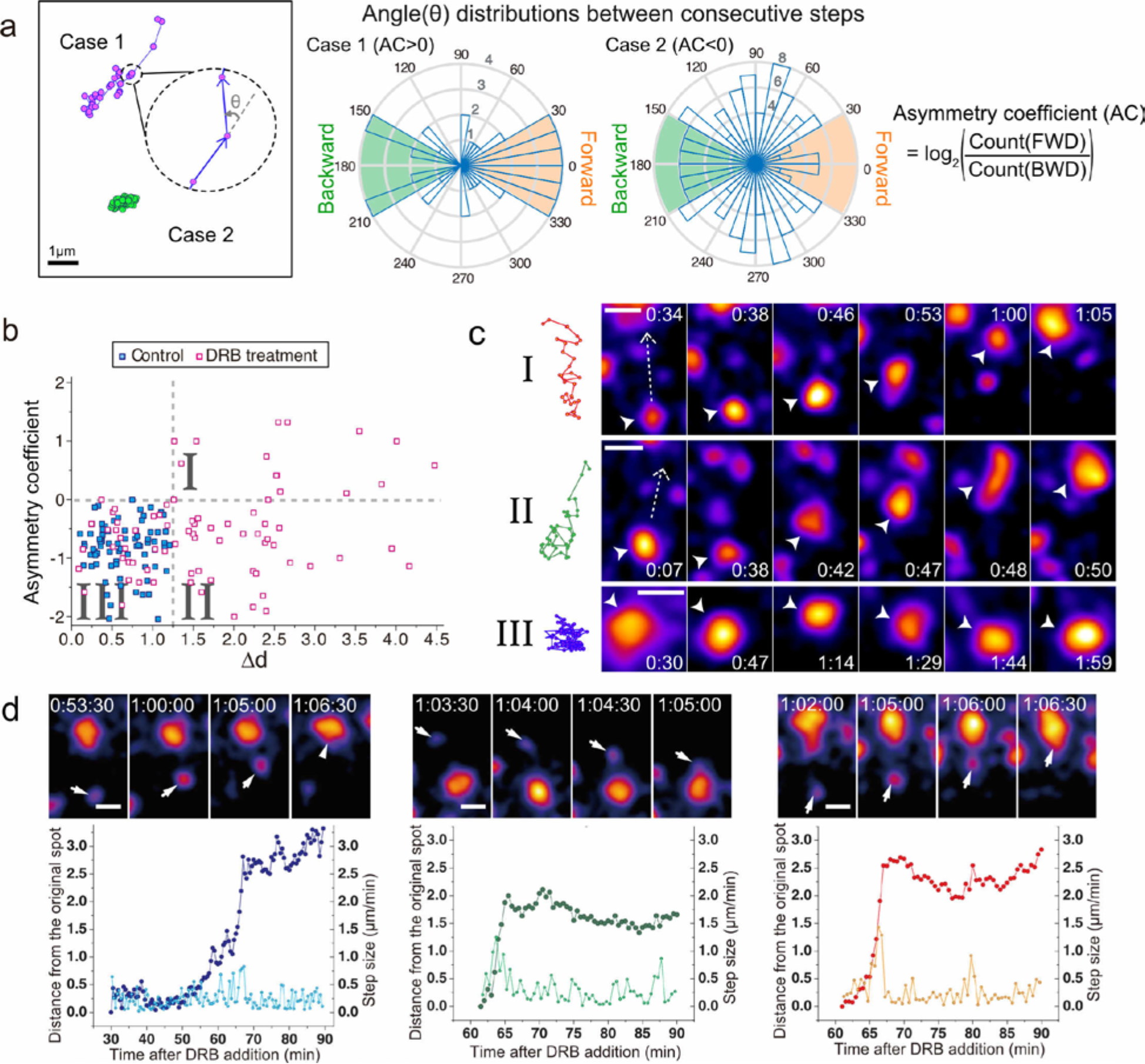
DRB treatment induces long-range, directional speckle movements. **a.** Analysis of angular distributions for two speckle trajectory examples: (Left) Two speckle trajectories- Case 1 (purple), Case 2 (green). The angle (θ) between every two consecutive steps is calculated (cartoon blowup). (Middle) Histogram distributions of angles for Case 1 (left) compared to Case 2 (right) trajectories. Radius of shaded sectors (blue) shows number of adjacent steps with a given angle. Backward (BWD) steps are defined by angles in green; forward (FWD) steps are defined by angles in orange. Asymmetry coefficients (AC) for Case 1 and Case 2 are negative and positive, respectively, as AC (right) is defined as the log2 ratio of FWD over BWD steps. **b.** Scatterplot of asymmetry coefficient (AC, y axis) versus net displacement (Δd, x axis) for speckle trajectories from control (blue squares) versus DRB-treated cells (pink squares). 3 trajectory types I-III are defined by their placement within three quadrants (I-III) defined by dotted lines (AC=0; Δd = 1.25). Control cells fall into negative AC and low Δd quadrant (I). n=75 speckles from 10 control cells; n=80 speckles from 10 DRB-treated cells. **c.** Speckle trajectories (left) and corresponding time-lapse images (right) from each of the 3 types I-III in (b). Time (hr:min) is after DRB addition. Scale bars: 1 μm. White dashed arrow marks direction of movement; arrowheads mark moving speckle. (Top) Type I: Forward motion-dominant, long-range directional motion; (Middle) Type II: long-rang directional motion, but fewer forward steps. (Bottom) Type III: Confined speckle motion. See Supplementary Video 1. **d.** Three examples of long-range directed speckle motion. (Top) Time-lapse images. Scale bars: 1 μm. White arrowheads mark speckle that moves. (Bottom) Distance (μm, vivid colors, left y-axis), measured from the starting spot position, and velocities (μm/min, light colors, right y-axis) as a function of time after DRB addition.

To compare speckle mobility in control versus DRB-treated cells, we created a scatter plot summarizing AC values versus net distance displacement for speckles (Fig. 2b). This scatter plot reveals three motion regimes (Fig. 2b, c): Type I trajectories show large net displacements (≧1.25 μm) with positive AC values as a result of subtrajectories that contain a large number of discrete forward steps. Type II trajectories show large net displacements but negative AC values. In contrast, Type III trajectories (Fig. 2b, c, Supplementary Video 1) show small net displacements with negative AC values, consistent with a confined motion of speckles.

In control cells, essentially all speckles showed locally confined, Type III trajectories. No speckles (0/75) traveled further than 1.25 μm, and AC values were all below 0, with a total AC range of ~ −2.0-0.0. In contrast, in DRB-treated cells, speckle mobility and AC values increased significantly; 63% of speckle trajectories (Type I+Type II, 50/80) exhibited net distance displacements, Δd, greater than 1.25 μm, with 14 out of 50 of these trajectories showing AC values greater than 0 (Type I, Fig. 2b). Several of these speckles moved as far as 3 - 4 μm, nearly three times as far as the largest displacement seen in control cells.

Plotting the distance moved as a function of time revealed that the subtrajectories containing periods of long-range movement were predominately unidirectional with larger steps than stationary movements shown before or after the long-range movement. (Fig. 2d). The imprecision of the spatial alignment of nuclei between time points, due to changes in nuclear shape, prevents us from determining to what degree small reverse speckle movements represent true reverse speckle movements versus alignment errors. Projecting over time the 2D spatial projections from multiple time points revealed the linear path of these subtrajectories containing long-range directional movements (Supplementary Fig. 2).

### Repeated long-range, directional speckle movements all terminate with speckle fusion

Strikingly, all the examples of long-range speckle movements shown in Fig. 2 and Supplementary Fig. 2 terminate with a fusion between the smaller, mobile speckle and a larger, relatively immobile speckle. These are not exceptional examples but rather the general rule observed for nearly all long-range movements contained within a single observation period.

Fig. 3 shows all long-range speckle movements in two representative nuclei as examples. Ten long-range speckle movements in nucleus 1 (Fig. 3a, Supplementary Video 2) end with fusion to 7 larger speckles; in three cases, two different small speckles move long-distance, and merge with the same larger speckle. Nine long-range speckle movements in nucleus 2 (Fig. 3B, Supplementary Video 3) end with fusion to 5 larger speckles; three smaller speckles merge with the same larger speckle in one case and two smaller speckles merge with the same larger speckle in two other cases. Thus 19/19 long-range speckle trajectories terminate with fusion to a larger, mostly immobile nuclear speckle.

**Figure 3.**
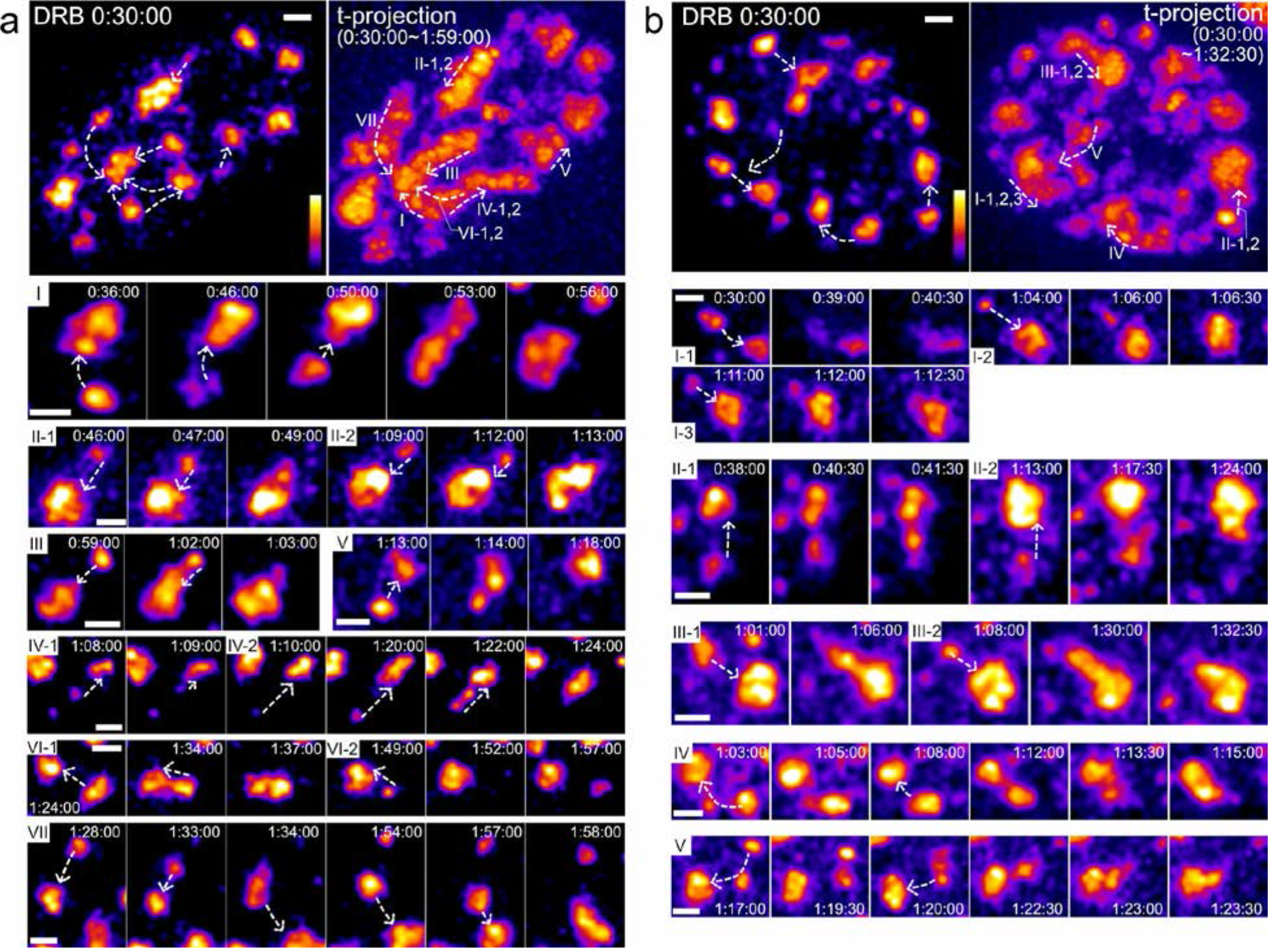
Long-range speckle movements are directed towards other speckles and terminate by speckle fusion. **a.** Overview of speckle motions in one nucleus (cell clone E8). All images show maximum-intensity 2D projections of optical sections, using pseudo-color intensity scale. Time (hr:min:sec) is after DRB addition. Scale bars: 1 μm. White dashed arrows show direction of speckle movements. (Top left) First time-lapse image taken 30 mins after DRB addition. (Top right) Maximum-intensity projection over all time-lapse 2D images. Roman numerals indicate different regions showing speckle movements, ordered by the time of speckle movements. Some regions show multiple speckles appear and then move. Each of these speckles is tagged by a number following the roman numeral (i.e. II-1,2 shows path of two different speckles moving in region II). (Bottom) Time-lapse images of each speckle movement marked by roman numeral and speckle number. See Supplementary Video 2. **b.** Same as (a) for second nucleus (cell clone D6). See Supplementary Video 3.

Again, projecting over time the 3D spatial projections from multiple time points revealed a largely curvilinear path for these long-range speckle trajectories, within the resolution of the nuclear alignment between time points (Fig. 3a,b, top right panels).

Even more strikingly, in many cases we saw repeated long-range speckle movements in which different nuclear speckles followed a very similar path to merge with the same single, larger speckle. In the two nuclei shown in Fig. 3, 6/12 larger speckles are the fusion targets for two or three mobile, smaller speckles. In each of these 6 examples, the second or even third smaller nuclear speckle moves along essentially the same linear path, within the approximate resolution of our nuclear alignment, to fuse with the larger speckle. Typically, after one speckle began to move on a long-range trajectory, a new speckle would nucleate and enlarge, either at the site of the first speckle prior to its movement (Fig. 4a,b, Supplementary Video 4) or along the trajectory followed by the first speckle (Fig. 4c,d, Supplementary Video 4, Supplementary Fig. 3). The second speckle would later begin to move along a similar trajectory to the first and then merge with the same speckle with which the first speckle moved.

**Figure 4.**
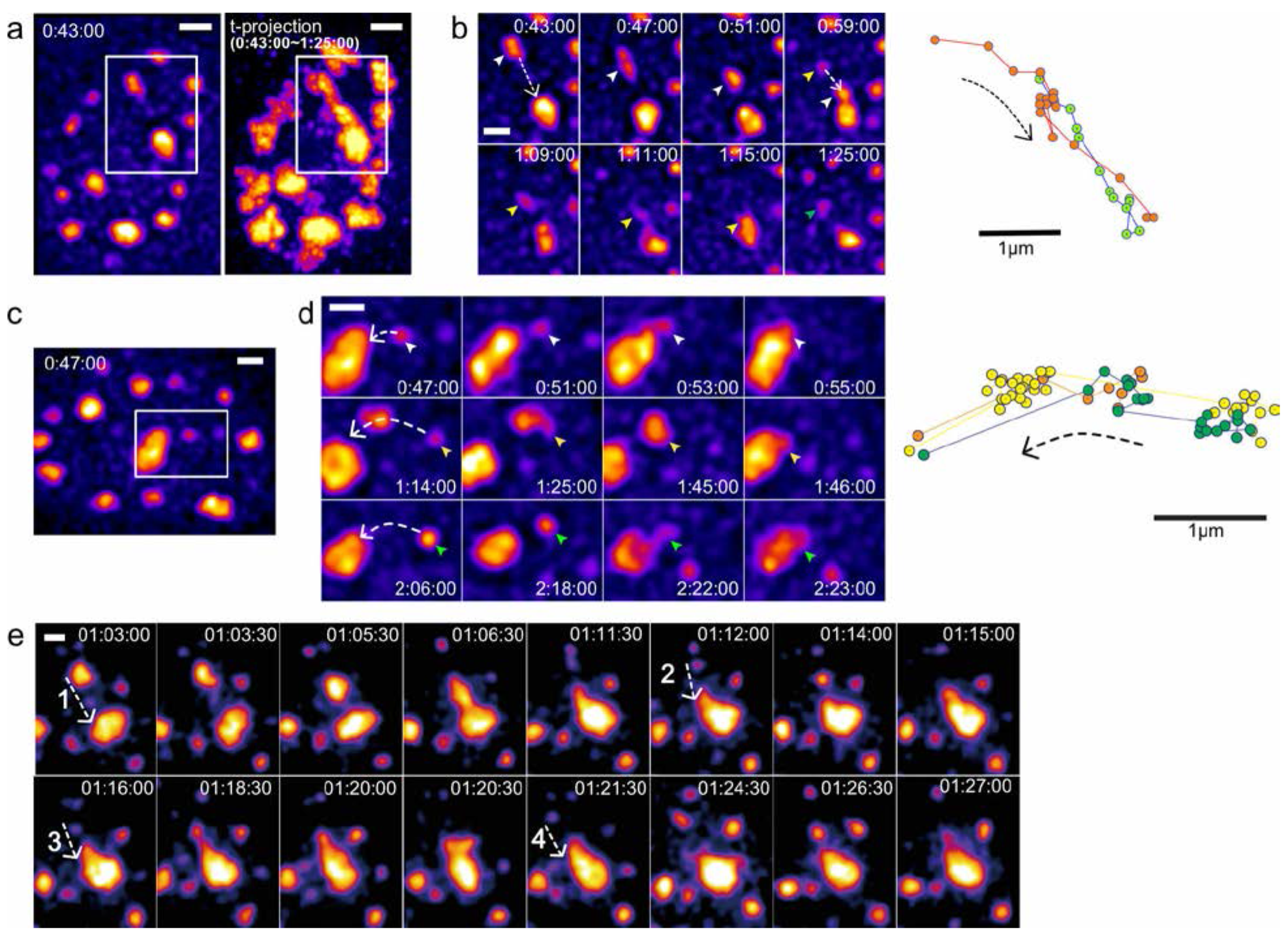
Multiple speckles undergo long-range, directional movements along similar paths: repeated cycles of speckle nucleation, motion, and fusion to same target speckle. 2D images represent 2D maximum-intensity projections of z-stacks. GFP-SON intensities are pseudocolored to increase dynamic range. Time (hr:min:sec) represents time after DRB addition. Scale bars: 1 μm. **a.** (Left) Image of first cell nucleus 43mins after DRB addition, with boxed region containing repeated long-range speckle motions shown in (b). (Right) Maximum-intensity projection of multiple 2D projected images over time from 43 mins to 1 hr 25 mins. An elongated high intensity “trail” is generated by the time projection of repeated speckle movements. **b.** (Left) Time-lapse images of speckle movements within boxed region in (a). White dotted arrow lines show the direction of speckle movements. First speckle (white arrowheads) moves and merges with target speckle. Second speckle (yellow arrowheads) appears along path followed by first speckle and moves to same target speckle. Green arrowhead marks appearance of third speckle along same path. (Right) Trajectories of first (brown) and second (green) moving speckles, showing their close overlap. **c.** Image of second cell nucleus 47 mins after DRB addition, with boxed region containing repeated speckle motions shown in (d). **d.** (Left) Time-lapse images of speckle movements within boxed region in (c). White dotted arrow lines show the direction of speckle movements. White, yellow, green arrowheads point to 3 different speckles that move over similar paths. (Right) Trajectories of these 3 speckles showing their close overlap. Second (yellow) and third speckles (green) nucleate at similar location but at different times. **e.** An additional example of 4 repeated speckle movements along similar paths. See Supplementary Video 4 for boxed regions shown in b, d, e.

We observed a number of cases in which three or even four speckles would appear in the same or similar location, observing long-range movements of 2, 3, or even 4 of these speckles along a similar trajectory before these speckles terminated their motion by fusion with the same larger speckle (Fig. 4e, Supplementary Video 4). Furthermore, we observed examples in which one speckle would merge with another, and then this merged speckle would move to fuse with a third speckle (Fig. 3a, VII, Fig. 3b, V, Fig. 4d, 1:14:00-1:46:00).

These observations of long-range movements occurring through largely unidirectional steps along curvilinear trajectories argue strongly for directional speckle movements along or within a nuclear path or channel. This is strongly supported further by these observations of repeated speckle movements along the same path or channel. Finally, as a further suggestion for a mechanism leading to directed movements along a possible path, we observed that speckles undergoing long-range movements would sometimes elongate in the direction of motion (Fig. 4b, 00:47:00, 1:09:00, Fig. 4e, 1:06:30) or the recipient speckle would form a long protrusion along the path of the original smaller speckle trajectory (Fig. 3a, I, 00:53:00, III, 1:02:00, Fig. 3b, III, 1:06:00, 1:30:00, Fig. 4d, 2:22:00, Fig. 4e, 1:18:30).

### Viscoelastic behaviors of speckles and estimation of speckle viscosity

During long-ranged speckle movements, we observed distinctive viscoelastic behaviors such as “inchworm” or reptation motion, suggesting speckles as liquid-like bodies. For example, a speckle would elongate in the direction of another speckle and then retract at the elongated end nearest the target speckle to from a rounded speckle now closer to the target speckle; this was then followed by a second round of elongation towards the target speckle followed by fusion and then rounding into a single, larger speckle (Fig. 5a). In other cases, one speckle would significantly elongate towards another target speckle until a long, linear speckle was formed that would fuse with the target speckle; this would be followed by retraction of the elongated speckle into the target speckle to form a single, rounded and larger speckle (Fig. 5b-d).

**Figure 5.**
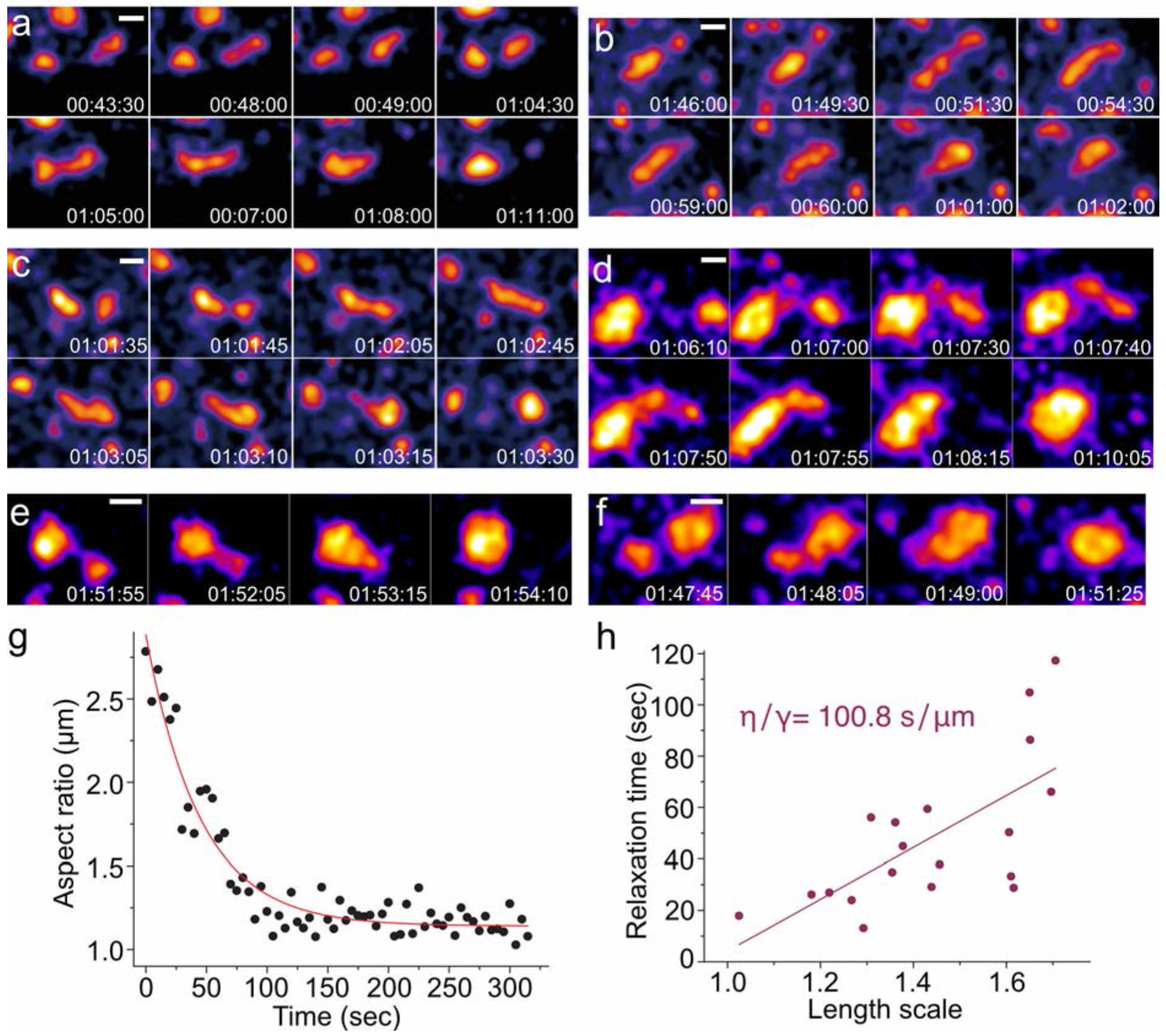
Viscoelastic behaviors of nuclear speckles. SON-GFP images represent maximum intensity 2D projections of 3D image stacks. Time (hr:min:sec) is after DRB treatment. Scale bars: 1 um. **a.** “Inch-worm“-like motion: A nuclear speckle repeats cycles of elongation and translocation until it fuses with another speckle located ^~^3 μm away from starting position of first speckle. **b-d.** reptation motion: A speckle elongates towards and then fuses with another speckle prior to speckle rounding. **e-f.** Two examples of most commonly observed speckle fusion. **g.** Plot of the aspect ratio versus time for two speckles fusing with each other. The Aspect Ratio is defined as the ratio of the long axis (l_long_) to short axis (l_short_) lengths of the ellipse approximating the morphology of the fusing speckles. Fitting this plot to a single exponential decay curve yields an exponential decay constant corresponding to the relaxation time of the speckle fusion. In this example, the relaxation time was calculated as 44.9 sec. **h.** Plot of relaxation time versus length scale for multiple speckle fusion events (N=19). The slope of this linear fit of relaxation time versus length scale, corresponds to the inverse capillary velocity, equal to the ratio of viscosity to surface tension (η/γ). The measured inverse capillary velocity from this plot = 100.8 s/μm.

Furthermore, examination of speckle fusion events (Fig. 5e, f) supports the concept of nuclear speckles as liquid-like nuclear bodies which have a similar viscosity as measured previously for other liquid-like RNP bodies such as nucleoli and P granules^10, 11, 30^. We estimated the viscosity of speckles by analyzing the time required after speckle fusion for a relaxation to a circular shape in cells treated with DRB. As expected, relaxation times (τ) were linearly proportional to the characteristic length scale (L) with a slope, which should approximate the inverse capillary velocity, of 101 s/μm (Fig. 5h). This value is similar to previously measured inverse capillary velocities of 2 s/μm for germline P granules of C. elegans^10^, 46.1 s/μm for nucleoli of *Xenopus* oocytes^11^, and 125 s/μm for large RNP assemblies (grPB) in C. elegans oocytes^30^. Assuming a speckle RNP granule size of ~20 nm, we obtained an estimated viscosity (η) of speckles of ~1·10^3^ Pa·s, close to the measured viscosities of nucleoli (~2·10^3^ Pa·s) and grPB (5·10^3^ Pa·s).

### Long-range speckle motion does not require polymerized actin

Previous examples of long-range chromosomal movements within interphase nuclei suggested a direct or indirect role of actin, and in certain cases nuclear myosin 1C, in these movements^31, 32^. To determine whether actin was involved in the long range motion of speckles, we inhibited actin polymerization using latrunculin A (latA), and then measured the frequency and length distributions of of long-range speckle movements.

LatA addition resulted in a decrease of long-range movements (Fig. 6a, Table 1). After latA addition, long-range speckle movements greater than 1 μm were observed an average of 2.7 times per nucleus. This compared to an average of 9.8 long-range speckle movements per nucleus in control cells. Although the frequency of long-range nuclear speckle movements was significantly reduced, the distance distribution of these long-range speckle movements did not significantly change with latA addition relative to those observed in control cells (Fig. 6a). Similar types of long-distance speckle trajectories, also ending in fusion with a target speckle, were observed after latA treatment (Fig. 6b, c, Supplementary Video 5).

**Figure 6.**
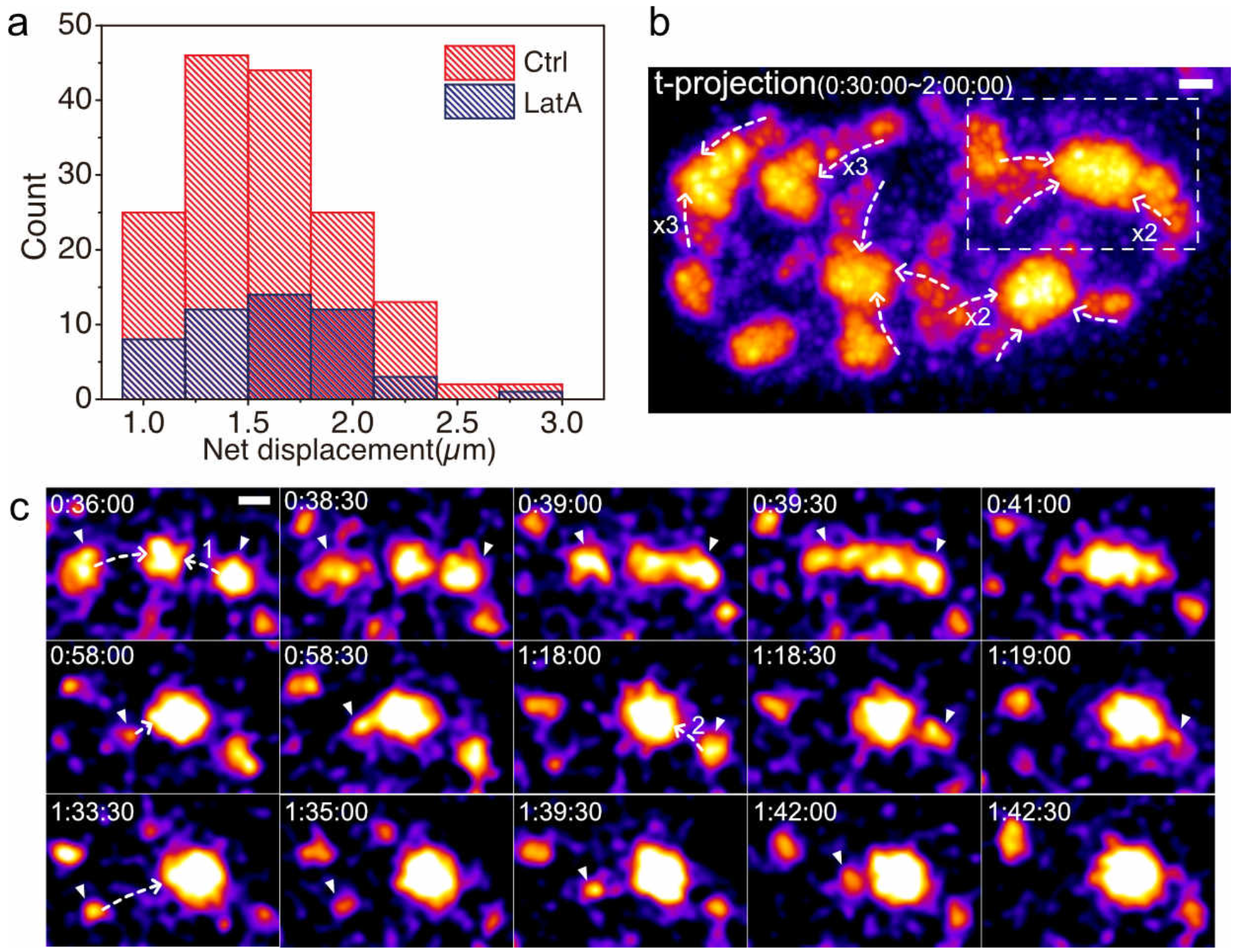
Long-range speckle motion does not require polymerized actin. **a.** Histogram of speckle net displacements after DRB addition with or without latA. LatA treatment decreases the frequency of long-range speckle motion, but not the length distribution of their net displacements. **b.** Maximum-intensity projection over time (t-projection) of GFP-SON nuclear 2D projections from 30 mins to 2 hrs after DRB and latA treatment. This t-projection shows curvilinear paths of long-range speckle motions which terminate at another speckle. White dotted lines indicate direction of speckle motions. Numbers show the number of speckles that move along the direction of the arrow over time (no number for one speckle). Time (hr:min:sec) represents time after DRB addition. Scale bar: 1 μm. See Supplementary Video 5. **c.** Time-lapse images from the boxed region in (b). Arrowheads point to moving speckles, and dotted lines indicate direction of speckle movements. The number on the dotted line distinguishes different speckles that move on the same path. Time (hr:min:sec) represents time after DRB addition. Scale bar: 1 μm.

**Table 1.** Numbers of long-range speckle movements (>1 μm) per nucleus in cells treated with latA or expressing an actin mutant after DRB addition. “T” stands for transfected cells with indicated actin construct, “U” stands for non-transfected control cells in same cell dish. LatA was added 30 mins prior to DRB addition. DRB was added at time 0, and time-lapse measurements were made from 30-90 mins after DRB treatment.

| | Number of cells | Average number of long-range movements per cell | Average distance($\mu\text{m}$) |
| --- | --- | --- | --- |
| Control | 16 | 9.8 | 1.53 |
| LatA | 18 | 2.7 | 1.50 |
| WT actin(T) | 17 | 13.4 | 1.48 |
| WT actin (U) | 17 | 12.8 | 1.48 |
| G13R actin(T) | 20 | 12.3 | 1.28 |
| G13R actin(U) | 15 | 12.2 | 1.28 |
| S14C actin(T) | 17 | 14 | 1.38 |
| S14C actin(U) | 22 | 13.5 | 1.34 |

To further test the possible involvement of actin in speckle motion, we transiently transfected cells with plasmid constructs of actin fused to mRFP and a nuclear localization signal (NLS) to concentrate this actin fusion protein in the nucleus^33, 34^ Different constructs were used for expression of the wild type beta-actin sequence, a nonpolymerizable actin mutant^34^ (G13R), or an actin mutant that favors polymerized actin F-actin^33^ (S14C).

Long-range nuclear speckle movements were observed in cells transfected (T) with all three mRFP-NLS-actin mutants or mRFP-NLS-wild-type actin as well as nontransfected control cells (U) (Table 1). The average number of long-range movements nuclear speckle movements ranged from ~12-14 per nucleus. Mean distances of these speckle movements were also similar (1.28-1.48 μm). The velocities of speckle movement during the periods of long-range movement were also similar (~0.5-1.5 μm/min, data not shown).

Thus, we conclude that the observed long-range, directional movements of nuclear speckles after DRB treatment are not actin-dependent.

### Nuclear speckles move in DAPI-depleted regions that are enriched with SON-granules

To explore a possible structural basis for the repeated directed movement of small nuclear speckles along the same apparent path towards a larger nuclear speckle, we used superresolution light microscopy. 3D structured illumination microscopy (SIM) and stimulated emission depletion (STED) microscopy on fixed cells was complemented with correlative livecell in which we used conventional wide-field microscopy on live cells followed by fixation and then SIM and STED imaging on the same cells.

Inspection of nuclear speckle movement in live-cell movies suggested the formation of transient, faint connections of GFP-SON between nearby speckles (Supplementary Videos 6, 7). Using STED microscopy, we could visualize individual, distinct SON-GFP granules that are in the size range of STED resolution of ~60 nm and are distributed nonrandomly in the nucleoplasm outside of nuclear speckles (Fig. 7). We used a combination of 3D SIM and STED to visualize the relative distribution of chromatin and SON. 3D SIM of DAPI-stained, fixed nuclei was followed by STED imaging of the same cells on a different STED microscope. These DAPI SIM and anti-GFP-SON STED images were then aligned using a cross-correlation procedure.

**Figure 7.**
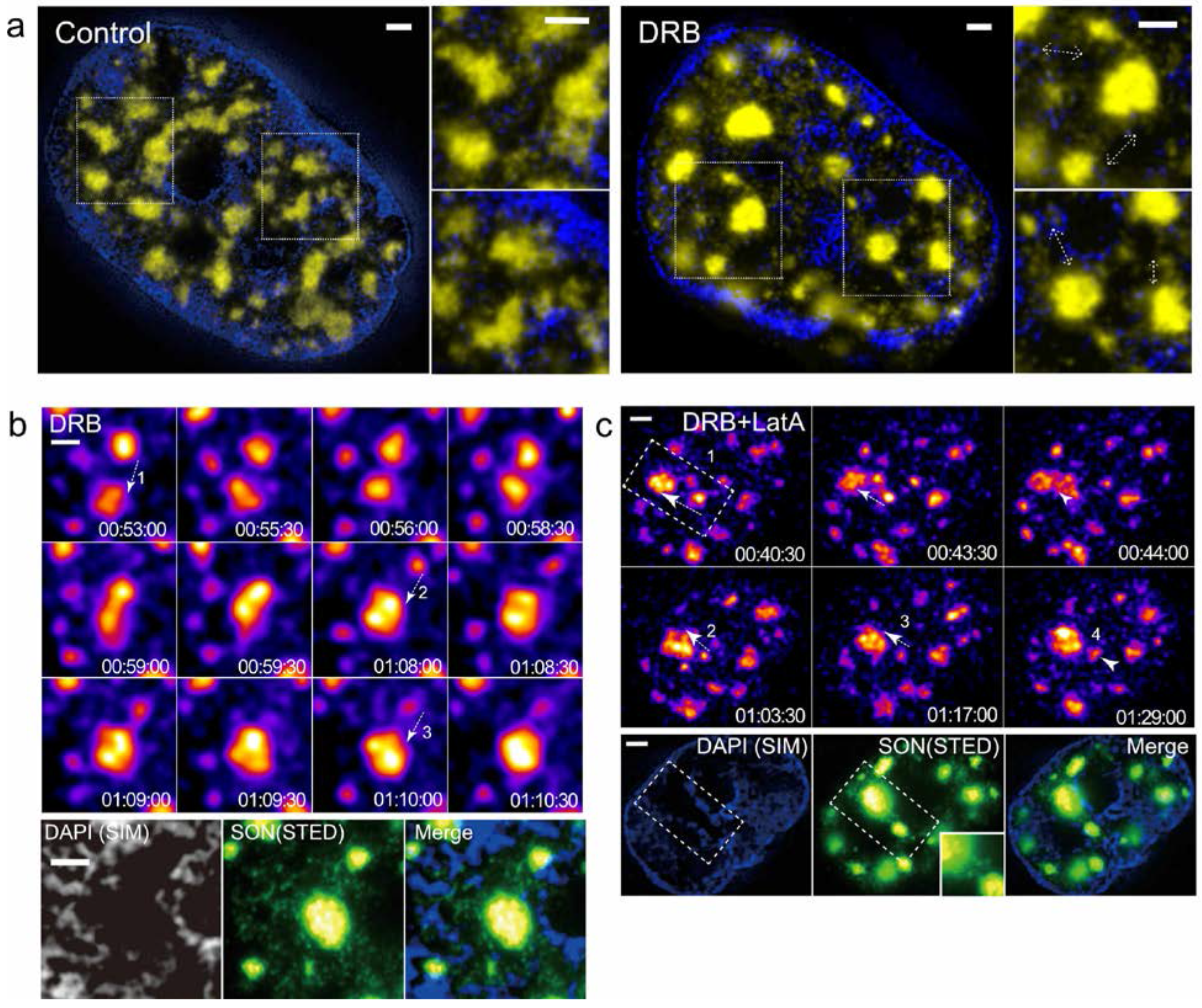
Repeated nuclear speckle motions occur within DAPI-depleted regions enriched in SON granules. Scale bars: 1 μm. **a.** Correlative SIM (DAPI, blue) and STED (SON, yellow) images of single optical sections, showing heterogeneous distribution of nuclear speckles and SON granules in DAPI-poor regions before (left) and after (right) DRB treatment. Regions of interest are marked with dotted boxes and shown enlarged in panels to right of nuclear images. Dotted arrows (right panels, right side) are drawn adjacent to local accumulations of SON granules running between two nearby nuclear speckles. **b.** Correlation between the path of repeated speckle motions, DAPI-depleted region, and local concentration of SON granules in DRB-treated cell. (Top) Time-lapse images of repeated speckle motions. Each speckle motion is marked by arrow and number. Time (hr:min:sec) is after DRB addition. See Supplementary Video 6. (Bottom) Correlative super-resolution images after fixation in region of repeated speckle movements(Top) occurred in DAPI-depleted region (left) enriched in SON granules (middle). Merged image (right). **c.** Same as (b) but for latA and DRB-treated cell. See Supplementary Video 7.

In these combined DAPI-SON images, we observed enrichment of SON granules in DAPI-depleted spaces connecting nearby nuclear speckles, with most speckles appearing to be connected by high local concentrations of these SON granules (Fig. 7a). These connections persisted even after DRB treatment, even becoming clearer due to the reduced number of granules overall in the nucleoplasm (Fig. 7a, right). The spatial distribution of these granules suggests they are related to the faint, transient GFP-SON connections between nuclear speckles visualized in our live-cell imaging.

We next used correlative live-cell and super-resolution imaging to determine the relationship between the path followed by repeated nuclear speckle trajectories and these DAPI-depleted, SON-granule-enriched spaces connecting neighboring nuclear speckles. After DRB treatment, we followed live-cell imaging with immediate fixation of cells using paraformaldehyde and staining. Inspection of the live-cell movies identified cells containing examples of repeated speckle movements along a similar path terminated by fusion with the same larger nuclear speckle. These cells were then identified and imaged by 3D SIM and STED.

These correlative images revealed that the path or channel along which nuclear speckles had repeatedly moved was relatively DAPI-depleted but surrounded by DAPI stained chromatin (Fig. 7b). Moreover, a relatively high concentration of SON granules was present within these paths or channels relative to surrounding nuclear regions. Similar correlative results were observed in cells treated with latA prior to addition of DRB (Fig. 7c).

Our observations of nuclear speckles moving within DAPI-depleted regions suggest that chromatin may act as a barrier to speckle motion. We next used live-cell imaging to examine the temporal correlation between changes in the DNA distribution surrounding nuclear speckles and speckle movement and fusion events (Fig. 8). We used the cell-permeable, far-red SiR-Hoechst dye that binds DNA in the minor grove like DAPI. Chromatin shows condensation with time after DRB treatment, increasing the size of DNA-depleted intranuclear spaces (Fig. 8a). Fusion between nearby large speckles indeed is temporally correlated with the disappearance of the DNA staining observed in preceding time points within the space separating these same speckles (Fig. 8b-e, Supplementary Videos 8, 9).

**Figure 8.**
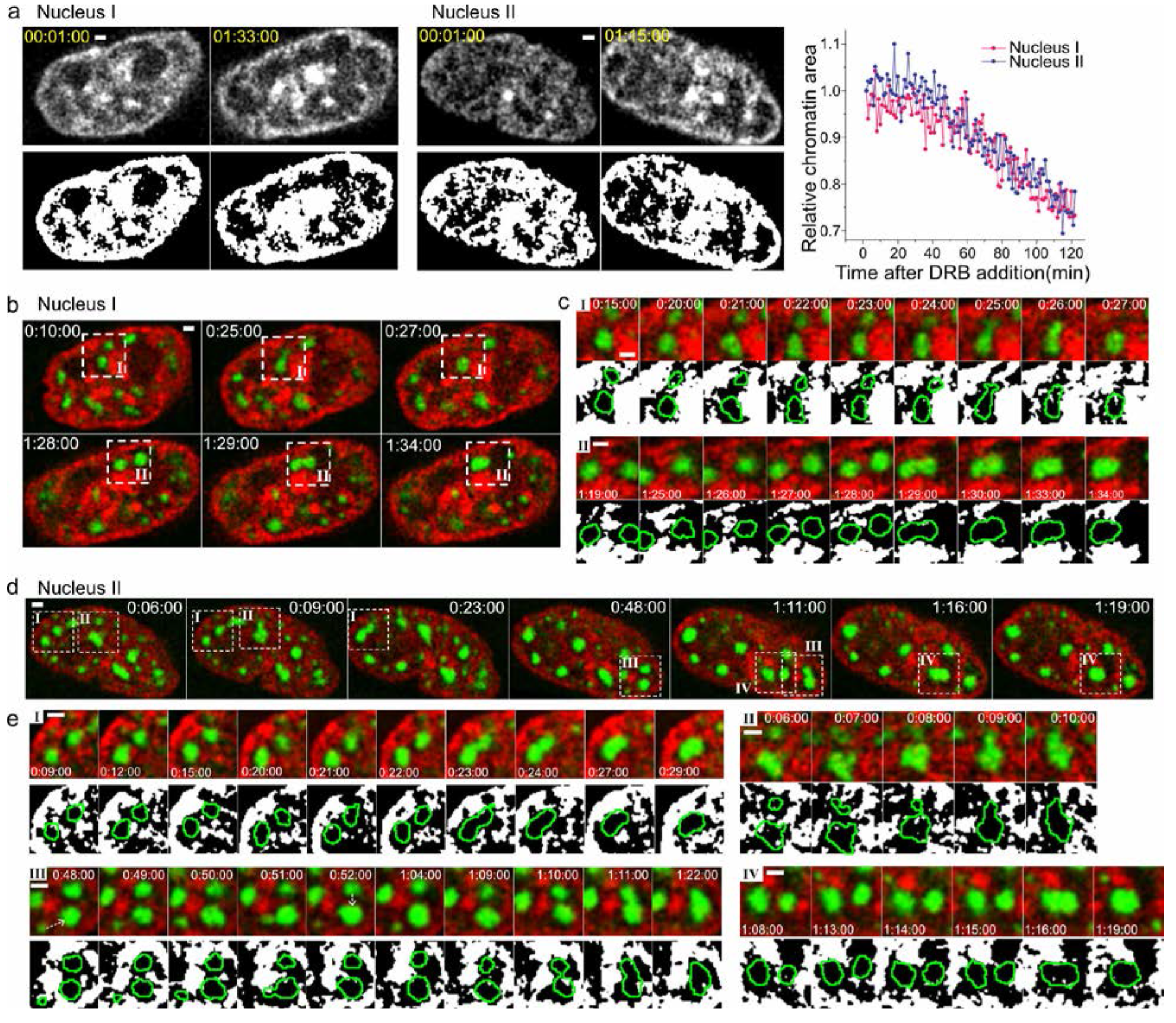
Speckle motions are spatially limited by chromatin structure. All images show single optical sections. Time (hr:min:sec) is after DRB addition. Scale bars: 1 μm. **a.** Chromatin compaction over time after DRB addition. (Left, Middle) Gray-scale (top) and binary (bottom) images of chromatin (SiR-Hoechst) in nucleus I (left) and nucleus II (middle) at indicated time points. (Right) Change in relative chromatin area (y) over time after DRB addition (x). **b.** Time-lapse images of nucleus I showing speckle (GFP-SON, green) motions and fusion following loss of chromatin (SiR-Hoechst, red) between the speckles that fuse. See Supplementary Video 8. Regions of interest (ROI) I and II marked by dashed lines are shown enlarged in (c). **c.** (Top row for ROI I and II) Time-lapse images show speckle-merging events occurred after depletion of chromatin in space between speckles. (Bottom row for ROI I and II) Binary images of thresholded SiR-Hoechst staining (white) relative to speckle boundaries (green lines). **d.** Same as (b) for nucleus II. See Supplementary Video 9. Nucleus II contains four ROI (I-IV). **e.** Same as (c) for nucleus II.

### Similar long-range and repeated speckle movements after heat-shock

To place our observations within a more physiological context, we examined speckle dynamics during heat shock. After heat shock, while most genes are transcriptionally down-regulated 5-fold or more (Mahat, 2016), heat-shock genes are induced up to 200-fold (Lis, 1981, Mahat, 2016). Previously, we demonstrated that heat-shock transcriptional induction of Hsp70 plasmid transgene arrays was greatly enhanced after nuclear speckle contact (Khanna et al. 2014). Similar to DRB treatment, after heat shock we observed multiple examples of long-range motions of smaller nuclear speckles terminating with fusion with larger nuclear speckles. Supplementary Fig. 4 shows a striking example of repeated nucleation of new speckles adjacent to the Hsp70 transgene array followed by long-range motion of these speckles towards a larger, distant nuclear speckle. Later, speckles nucleate along this speckle trajectory and then merge to form a continuous, ~ 4 μm long connection between the transgene array and distant speckle. Supplementary Fig. 5 shows a long-range speckle motion terminating after association with the Hsp70 BAC transgene array which then shows an abrupt increase in its transcription.

## Discussion

Here we showed an increased mobility of nuclear speckles after transcriptional inhibition by DRB, including prominent long-range movements of smaller speckles over distances of 1-4 um with velocities 0.2-1.5um/min that typically terminated through fusion with a larger speckle. These long-range movements appeared directional, following roughly linear trajectories pointing to and ending with merging in pre-existing stationary speckles. Speckles frequently elongated in the direction of movement during these movements, suggesting the possibility of an active transport mechanism. Directionality was also inferred based on our observations of repeated cycles of nucleation of new speckles and their movement along similar trajectories followed by previous moving speckles. These long-range, speckle movements occurred independent of polymerized nuclear actin as their length distribution was unchanged after lantrunculin A treatment. Correlation of live-cell imaging with super-resolution microscopy on fixed samples revealed that speckles moved within channels relatively depleted of DNA but enriched in SON-protein containing granules.

As observed previously, upon transcriptional inhibition speckles became rounder and larger. This increase in speckle size was imagined to occur as a result of RNA processing components being released from previous sites of transcription, diffusing into speckles, and then accumulating within the speckle through binding to other components^23, 35^. Dephosphorylation of SR proteins was proposed to lead to their accumulation in speckles, while phosphorylation of SR proteins was proposed to lead to their release from speckles, diffusion into the nucleoplasm, and binding to local transcription sites^36, 37^. Consistent with this model, modeling the mobility of speckle proteins as measured by FRAP (Fluorescence Recovery After Photobleaching) led to the conclusion that differential nuclear distribution of splicing factors was controlled by changes in
their binding to either nucleoplasmic or speckle sites^38, 39, 40^.

Not explained by these studies was how the number of speckles decreases with transcriptional inhibition. Also these previous studies proposed that the increased size of speckles after transcriptional inhibition is entirely due to diffusion of speckle components from the nucleoplasm followed by binding to sites within speckles.

Our results instead suggest that a significant cause of both the increase in individual speckle size and brightness and the reduction in speckle number is through the directed, long-range movements of speckles towards other speckles followed by their fusion. Future work will be needed to determine the relative contributions of diffusion and binding of speckle components versus speckle fusion events to the observed increase in speckle size observed after transcriptional inhibition. Future work also will be needed to determine whether posttranslational modifications of speckle proteins are also connected to directional motion and/or fusion of speckles.

During the dynamic movement of speckles, we confirmed liquid-state properties of nuclear speckles. Speckles exhibited nucleation, deformation in the direction of translocation, and fusion. Strikingly, we found that motion of speckles after transcriptional inhibition always ended with fusion with another speckle. After fusion, speckles regained their circular shape but were bigger and showed an increased intensity, suggesting mixing of constituent molecules as seen in liquid-phase cellular bodies^41^. Analysis of the time interval between speckle fusion and rounding of the merged speckles allowed us to estimate the viscosity of nuclear speckles, which we found was similar to previously measured viscosities for *Xenopus* oocyte nucleoli and grPBs of *C. elegans* oocytes. Not considered in our analysis was the possibility that adjacent chromatin interacting with nuclear speckles might slow the rounding of the nuclear speckle after fusion events. Therefore, our estimates of nuclear speckle viscosity might be best considered an upper bound on the actual speckle viscosity, where the actual viscosity nuclear speckles might be somewhat lower depending on how significant these chromatin interactions might be on the actual relaxation time of the speckles after fusion. Overall, though, these measurements indicate that nuclear speckles possess viscoelastic properties and behaviors similar to those observed in other RNP bodies.

Significantly, although the frequency of long-range speckle movements decreased after perturbation of actin polymerization, the length distributions of these speckle movements did not change. Therefore, the long-range, directed movement of speckles after transcriptional inhibition is not dependent on polymerized actin. This rules out a more conventional actin-myosin based mechanism for speckle motility, despite reports of nuclear myosin I isoforms in nuclear speckles and enrichment of these isoforms in speckles after transcription inhibition^42^.

Fusion between nearby speckles to form larger speckles could be explained as driven by collisions between speckles mediated through random Brownian motion. However, our observation of directed motion of speckles over distances up to several microns in length, in many cases followed by a new speckle moving along a similar path, requires a different explanation than random diffusion.

One possible clue pointing to a potential mechanism underlying speckle long-range motion might be our observation of SON-containing granules concentrated in DAPI-poor channels connecting nearby speckles. Correlative live-cell / fixed-cell imaging revealed that these channels roughly co-localized to the path of actual speckle movements observed in live-cells shortly before fixation. We can envision more than one scenario that might give rise to these localized SON-granule accumulations on the path of speckle movements.

A flow of SON-granules between neighboring speckles, through some unknown mechanism(s), might establish this concentration of granules. Numerical modeling of a simple binary fluid mixture predicts a net flux of individual components through diffusion from small to larger droplets due to the greater stability of the larger droplets because of their lower interface free energy. Moreover, this net flux generates a composition correlation between two neighboring droplets of different sizes, leading to an intradroplet gradient of interfacial tension and producing a hydrodynamic flow of small droplets towards larger droplets followed by droplet fusion^43^. Interestingly, this modeled behavior of droplet movements in simple binary fluid mixtures mirrors our observation of small speckles moving towards larger speckles. Additionally, movement of one speckle along a trajectory might leave SON-granules along its path-analogous to tiny droplets of water left behind by movement of a large water droplet along a surface. These pre-positioned SON-granules speckles might work as “steps” biasing movements of other speckles at later times. Finally, we cannot rule out an active, but unknown transport mechanism driving speckle movements. Additional experiments including future live-cell imaging at higher temporal and spatial resolution will be needed to explore these ideas.

Our correlative imaging using SIM, STED and live-cell imaging also allowed us to determine the relative spatial arrangement between chromatins and speckles. The movement of speckles occurred within the interchromatin space, with chromatin seen surrounding the speckles and the apparent channels through which the speckles moved, suggesting that chromatin may act as a spatial barrier to speckle movement and confine these movements to linear trajectories. Indeed, under normal growth conditions, we observed speckle mobility consistent with confined motion. The observed directional motion after transcriptional inhibition is consistent with a release of constraints to speckle motion as a result of the chromatin condensation observed in live-cell movies after transcriptional inhibition. Thus, we suspect that chromatin in nuclei plays a similar constraining role in restricting translocation and fusion of nuclear speckles as actin networks have been shown to play in constraining movement and coalescence of ribonucleoprotein droplets and nucleoli in oocytes^11, 44^.

In conclusion, in this study, the most striking aspects of the long-range speckle movement we observed were the repeated cycles of speckle nucleation, translocation of the newly nucleated speckles along a similar trajectory as that followed by preceding speckles, and then fusion with the same target speckle. Together these observations point to a previously unsuspected cellular trafficking system for movement and nuclear positioning of nuclear speckles. We anticipate these non-random speckle movements may provide a rapid and effective transport of RNA processing and/or transcription factors to specific nuclear sites. Moreover, recent identification of genomic regions that associate at near 100% frequencies with nuclear speckles^45^ suggest that the nuclear positioning of speckles may also drive nuclear genome organization. The phenomenology of nuclear speckle movements described here should allow future development of assays that can be used to identify the molecular components of responsible for this nuclear speckle cellular trafficking system.

## Materials and methods

### Cell culture and establishment of cell line

CHOK cells were grown in F12 media (Cell media facility, University of Illinois at Urbana-Champaign) supplemented with 10% Fetal Bovine Serum (Sigma-Aldrich, F2442) at 37°C in 5% CO_2_. To generate a stable cell line expressing EGFP fused to the SON protein, we used a BAC EGFP-SON transgene (EGFP-SON-Zeo BAC) in which a starting BAC containing a human SON genomic insert (RP11-165J2, Invitrogen) was retrofitted using BAC recombineering to add a Zeocin selectable marker and the EGFP sequence in frame with the SON NH2 terminus (Khanna, Hu, and Belmont 2014). We purified the EGFP-SON-Zeo BAC using the Large-Construct Kit (Qiagen). 5ug of the purified construct was linearized with BsiWI(NEB), followed by incubation at 65◻ for 20 mins to inactivate BsiWI. CHOK cells were transfected with the linearized EGFP-SON-Zeo BAC using Lipofectamine 2000 (Invitrogen) one day after passaging while at ~60% confluency following the manufacturer’s suggested protocol. After 36 hours, transfected cells were trypsinized (Trypsin-EDTA 0.25%, Gibco) and transferred to a larger flask to which selection media (200 μg/ml Zeocin, ThermoFisher) was added. After 10 days of selection, cells were subcloned by serial dilution into 96-well plates. Individual subclones were screened by microscopy using a Deltavision wide-field microscope (GE Healthcare) to select clones which showed uniform and stable EGFP-SON expression.

### Drug treatments

To add drugs without adding new serum, we removed half the media from the cell culture dishes, added chemicals to a 2x concentration using this media, and then returned this media back to the same dishes. The final DRB working concentration was 50μg/ml (Sigma, 50mg/ml stock solution in DMSO). Other drugs were used at the following working (and stock) concentrations: 50ug/ml α-Amanintin (A2263, Sigma, 1mg/ml in DI water), 0.1ug/ml triptolide (T3652, Sigma, 1mg/ml in DMSO), 200uM Cd solution (20920, Sigma, 100mM in DI water). For live cell imaging, chemicals were prepared in the same way, and returned to the live cell dish on the microscope.

For latrunculin A (latA) treatment (L5163, Sigma), cells were seeded on poly-L-lysine coated coverslips or glass-bottom dishes (Mattek). Coverslips or glass-bottom dishes (Mattek) were covered with 0.01% Poly-L-lysine solution (P4707, Sigma) for 10 mins, rinsed with sterilized DI water, and dried in the tissue culture hood. Cells were grown two days on these coated surfaces, treated with 1uM latA for 30 mins, and then with DRB as described above.

### Sample preparation for fixed cell imaging

For quantitative analysis of speckle morphology, cells were seeded on coverslips (Fisher) and fixed 48 hrs later at ~90-100% confluency using freshly prepared 3.6% paraformaldehyde (PFA, Sigma, P6148-500G) in PBS for 20 mins at room temperature. After three times of washing for 5minutes each in phosphate-buffered saline (PBS), the fixed cells were then mounted in a Mowiol-DABCO anti-fade medium^46^.

For STED and SIM imaging, immunofluorescence and DAPI staining were done after fixation and 3x 5min washes in PBS. For immunofluorescence staining, cells were incubated in blocking buffer (0.5 % Triton X-100 (Thermo Fisher) and 0.5 % normal goat serum (Sigma-Aldrich) in PBS for 30 mins. Cells were then incubated with monoclonal anti-SON primary antibodies (1:300 in PBS; Sigma-Aldrich) overnight at 4°C. After 3x 5min washes in PBS, cells were incubated with Atto647N-conjugated goat anti-rabbit IgG (1:300 in PBS; ATTO-TEC) for 2 hrs at room temperature, and then washed 3x for 5mins each in PBS. The cells were postfixed with 3.6% PFA for 15 mins, and washed 3x 5min in PBS. Cells were counterstained with 1ug/ml DAPI (Sigma-Aldrich) in PBST (0.1% Triton X-100 in PBS) for 0.5-1 hr at room temperature (Smeets et al. 2014), and then washed 5x for 5 mins each in PBS. We equilibrated the cell samples with Vectashield antifade mounting medium (H-1000, Vector Laboratories) for 10 mins, aspirated the media, and added fresh mounting medium. This was repeated 3-5times to completely infiltrate the cells with the mounting medium to avoid refractive index changes over the cells. Coverslips were mounted on slides and sealed with nail polish.

### Microscopy of fixed samples and analysis of cell morphology

We used a Deltavision wide-field microscope (GE Healthcare), equipped with a Xenon lamp, 60X, 1.4 NA oil immersion objective (Olympus) and CoolSNAP HQ CCD camera (Roper Scientific). 1024×1024 pixel 2D images were acquired as 3D z-stacks and processed using the “Enhanced” version of the iterative, nonlinear deconvolution algorithm provided by the Softworx software^47^ (GE Healthcare). 3D deconvolved image stacks were projected into 2D using a maximum intensity algorithm.

Analysis of speckle morphology used custom Matlab codes for image segmentation in several sequential steps. Multiple intensity thresholds and image segmentation steps allowed us to segment speckles of varying sizes and intensity levels. We first automatically choose an initial intensity threshold using Matlab’s *graythresh* function (*graythresh*\*1.5~1.8) based on Otsu’s method (Otsu, 1979). This allowed us to produce a binary image from which we measured area, perimeter, and form factor (FF= 4*π*Area/(Perimeter)^2^) of the larger, brighter speckles. To calculate intensities of these speckles, we used the segmented areas as a mask which was applied to a new 2D projected image generated by an intensity projection from the original 3D raw image stacks. The normalized speckle intensity was calculated by summing the intensities of all pixels within a speckle, subtracting the background intensity over an equivalent area and then normalizing by the speckle area ({Sum of pixel values in speckle-(mean value of nuclear background*size of the speckle)}/Size of speckle). Once these large speckles were segmented and analyzed, we removed them from the deconvolved projected image by replacing their pixel intensities equal to the cellular background level. We then repeated the image thresholding, setting a new intensity threshold using the *graythresh* function (*graythresh*\**0.8*~*1.8*), and repeating the same process as described above to measure a new set of speckles and then remove them from the image. If necessary, this process was repeated one more time to select the remaining small speckles. To prevent the situation in which part of a speckle that was not segmented in a previous cycle was counted as a new speckle, at each step we recorded the coordinates of segmented speckle centers and discarded any new speckles in a subsequent cycle that were within 0.3 μm of a previous speckle center. Through 3 such segmentation cycles, we were able to identify and measure the morphology of most speckles present in the nucleus. However, very small speckles less than 0.4 μm diameter were not considered in our measurements of changes in speckle morphology before and after transcriptional inhibition.

### Live cell imaging and tracking

For live cell imaging, we used a V3 OMX (GE healthcare) microscope, equipped with a 100X, 1.4 NA oil immersion objective (Olympus), two Evolve EMCCDs (Photometrics), a live-cell incubator chamber for CO_2_ perfusion, and two temperature controllers for the incubator and the objective lens heaters. Temperatures of the live cell chamber and lens were maintained at 37◻. Cells were seeded in glass-bottom dishes (Mattek) to reach ~90-100% confluency 48 hrs later. 3D stacks (z-spacing 300nm) were acquired in the conventional wide-field imaging mode at given time intervals, followed for each time point by 3D deconvolution and 2D maximum intensity projection using the Softworx software. Image J was used first to smooth these projections with a Gaussian filter (σ=2). Then a rigid body registration (ImageJ plugin ‘StackReg’) was applied to correct for any x-y nuclear rotation and/or translational displacement between sequential time points. A region including a full speckle motion was cropped from the image manually using ImageJ, and the single speckle was tracked by custom Matlab code. The cropped image was transformed into a binary image by the intensity threshold selected by Matlab’s *graythresh* function. The center of mass of the speckle was then determined over time by Matlab’s *regionprops* function and used for speckle tracking.

### Viscosity analysis

To estimate nuclear speckle viscosity, we acquired live cell movies using a 5 sec time interval over 90 mins. We identified nuclear speckles undergoing fusion and measured their long axes (l_long_) and short axes (l_short_) to calculate as a function of time their aspect ratio (AR), defined as AR=l_long_/l_short_. A relaxation time (*τ*) for the speckle fusion was estimated by fitting the measured AR to the exponential decay curve defined by AR= P+(AR_0_-P)·e^−t/τ^, where P is the plateau of the exponential curve and AR_0_ is the AR at time=0.

The length scale was calculated as L =[(l_long_ − l_short_)· l_short_]^0.5^ at time=0, and plotted against *τ*. From a linear fit of *τ* vs. L by least-squares estimation, we obtained the slope that corresponds to the inverse capillary velocity (η/γ), where η is the viscosity and γ the surface tension^48^. We could estimate γ ≈ k_b_T/d^2 49^, where k_b_ is the Boltzmann constant, T is the temperature, and d is molecular length scale (~10nm used for RNAPs in previous studies^11, 30^). We used ~20 nm for the characteristic RNP granules contained within IGCs^15, 16^. From the estimated γ and the inverse capillary velocity (η/γ), we estimated the viscosity of nuclear speckles.

### Correlative live cell imaging and super-resolution imaging

To combine live-cell imaging with STED and SIM, cells were seeded in glass-bottom dishes engraved with 50 μm grid pattern, letters and numbers (81148, Ibidi), reaching ~60-70% confluency 2 days later. Prior to live cell imaging, we recorded locations of target cells using the alphanumeric characters to find the same target cells later when we used different microscopes. After live-cell imaging, cells were immediately fixed by adding 2x PFA solution in PBS to the cell media. The final PFA concentration was 4%. Immunofluorescence and DAPI staining followed by mounting in Vectashield anti-fade medium were done as described above.

For STED images, we used a custom made STED microscope^50^, and acquired 3D z-stacks through the entire SON immunofluorescent stained nucleus using 300 nm z-spacing.

3D-SIM images were acquired on the V3 OMX microscope (see above) using 0.125 um z-spacing and sequential excitation at 488 and 405 nm on each image plane. Each z-stack image contained 15 images at 5 different phases per angle and 3 different angles per slice. We collected a full 3D-SIM image of GFP-SON and DAPI-staining. 3D-SIM image stacks were reconstructed with Softworx. The chromatic aberration offset between the GFP and DAPI wavelengths was measured with the alignment slide provided by GE Healthcare and used to correct the 3D-SIM images using the OMX Image registration function in Softworx.

To align the STED SON and SIM DAPI images, we selected a single STED optical z-section showing speckles of interest and then matched it to the most similar optical z-section from the complete stack of the 3D SIM GFP-SON image. Since 2D STED imaging was done using a 300 nm z-interval (z resolution ≈ 600 nm), we used the 2D projection of 3 adjacent SIM z-sections, consisting of the most similar SIM optical section plus the SIM optical sections immediately above and below this optical section. Because ‘StackReg’ registration tool in ImageJ required a combined stack of images, we matched x-y sizes of SIM image (1024×1024 pixels, pixel size= 40nm after reconstruction) and STED image (varying, pixel size=20~40nm) by adding pixels of background intensity or by taking a certain size of sub-region from a large image. We then generated a RGB SIM image from DAPI (blue) and SON (green) after saturating SON intensity to be used as a reference for alignment with STED image. We finally combined RGB SIM image and STED SON as z-stacks, and ran ‘StegReg’ with rigid body mode. In this way, STED SON image could be easily aligned on the strong SON signal of the SIM image, where SIM DAPI image and SIM SON image were grouped during alignment process. From the aligned z-stacks, we took SIM DAPI image and STED SON image.

### Live cell imaging and analysis of chromatin compaction after DRB treatment

To visualize chromatin structure in live cells, we incubated cells growing in glass-bottom dishes with 100 nM SiR-Hoechst (Spirochrome, SC007) for 1 hr before imaging. We started 3D live-cell imaging immediately after adding DRB, acquiring simultaneously both GFP-SON (excitation at 488nm) and SiR-Hoescht (excitation at 642nm) images for 2 hrs using a 1 min time interval. Deconvolution and maximum intensity 2D projection at each time point were performed by Softworx as described previously. We converted these time-lapse 2D projected images into binary images by segmentation using ImageJ plugin ‘Threshold’ based on the Otsu thresholding method^51^. From these segmented binary chromatin images, we computed the chromatin area as a function of time using the Matlab’s *‘regionprops’* function.

## Acknowledgements

This work was supported by National Institutes of Health grant R01 GM058460 to A.S.B, National Science Foundation grant PHY-1430124 to T.H., and National Institutes of Health Grant GM112659 to T.H.. T.H. is an investigator with the Howard Hughes Medical Institute.

## Author contributions

J.K. and A.S.B. conceived and designed experiments. J.K. conducted experiments, analyzed data. K.Y.H. acquired STED images, contributed on data analysis and experimental design. N.K. contributed to generate cell lines. A.S.B. and T.H. supervised the project. J.K. and A.S.B. wrote the paper with feedback from other authors.

## Competing interests

All authors declare no competing financial interests.

## Supplementary Videos

**Video 1. Speckle movements from trajectory types I-III in live CHO cell after DRB treatment.**

Corresponds to Figure 2C. Movie represents maximum intensity 2D projection of 3D image stack for each time point. Time (hr:min) after DRB addition and scale bar(1μm) are stamped on the movie.

**Video 2. Long-range movement and fusion of nuclear speckles in live CHO cell after DRB treatment.**

Corresponds to Figure 3A. Movie represents maximum intensity 2D projection of 3D image stack for each time point. Time (hr:min:sec) after DRB addition and scale bar are stamped on the movie.

**Video 3. Long-range movement and fusion of nuclear speckles in live CHO cell after DRB treatment.**

Corresponds to Figure 3B. Movie represents maximum intensity 2D projection of 3D image stack for each time point. Time (hr:min:sec) after DRB addition and scale bar are stamped on the movie.

**Video 4. Repeated long-range directional speckle movements after DRB treatment.**

Corresponds to Figure 4b, d and e. Each example was found in different cell nucleus. Movie represents maximum intensity 2D projection of 3D image stack for each time point. Time (hr:min:sec) after DRB addition and scale bar(1μm) are stamped on the movie.

**Video 5. Long-range movement and fusion of nuclear speckles after latrunculin A and DRB treatment.**

Corresponds to Figure 5. Movie represents maximum intensity 2D projection of 3D image stack for each time point. Time(hr:min:sec) after DRB addition and scale bar are stamped on the movies.

**Video 6. Repeated speckle movements along similar path and fusions with same target speckle after DRB treatment.**

Corresponds to Figure 6B. The local region of interest is marked with box in movie. Correlative STED and SIM images of SON and DNA (DAPI) are shown in Figure 6B following fixation and staining. Movie represents maximum intensity 2D projection of 3D image stack for each time point. Time(hr:min:sec) after DRB addition and scale bar are stamped on the movies.

**Video 7. Repeated speckle motions along similar path and fusions with same target speckle after latrunculin A and DRB treatment.**

Corresponds to Figure 6C. The local region of interest is in the center of the nucleus. Correlative STED and SIM images of SON and DNA (DAPI) are shown in Figure 6C following fixation and staining. Movie represents maximum intensity 2D projection of 3D image stack for each time point. Time(hr:min:sec) after DRB addition and scale bar are stamped on the movies.

**Video 8. Speckles fuse after chromatin barrier is depleted.**

Corresponds to nucleus I in Figure 7. SiR-Hoechst staining (red) with GFP-SON (green) after DRB treatment. Movie is a single optical section for each time point. Time(hr:min:sec) after DRB addition and scale bar are stamped on the movies.

**Video 9. Speckles fuse after chromatin barrier is depleted.**

Corresponds to nucleus II in Figure 7. SiR-Hoechst staining (red) with GFP-SON (green) after DRB treatment. Movie is a single optical section for each time point. Time(hr:min:sec) after DRB addition and scale bar are stamped on the movies.

**Supplementary Figure. 1.**
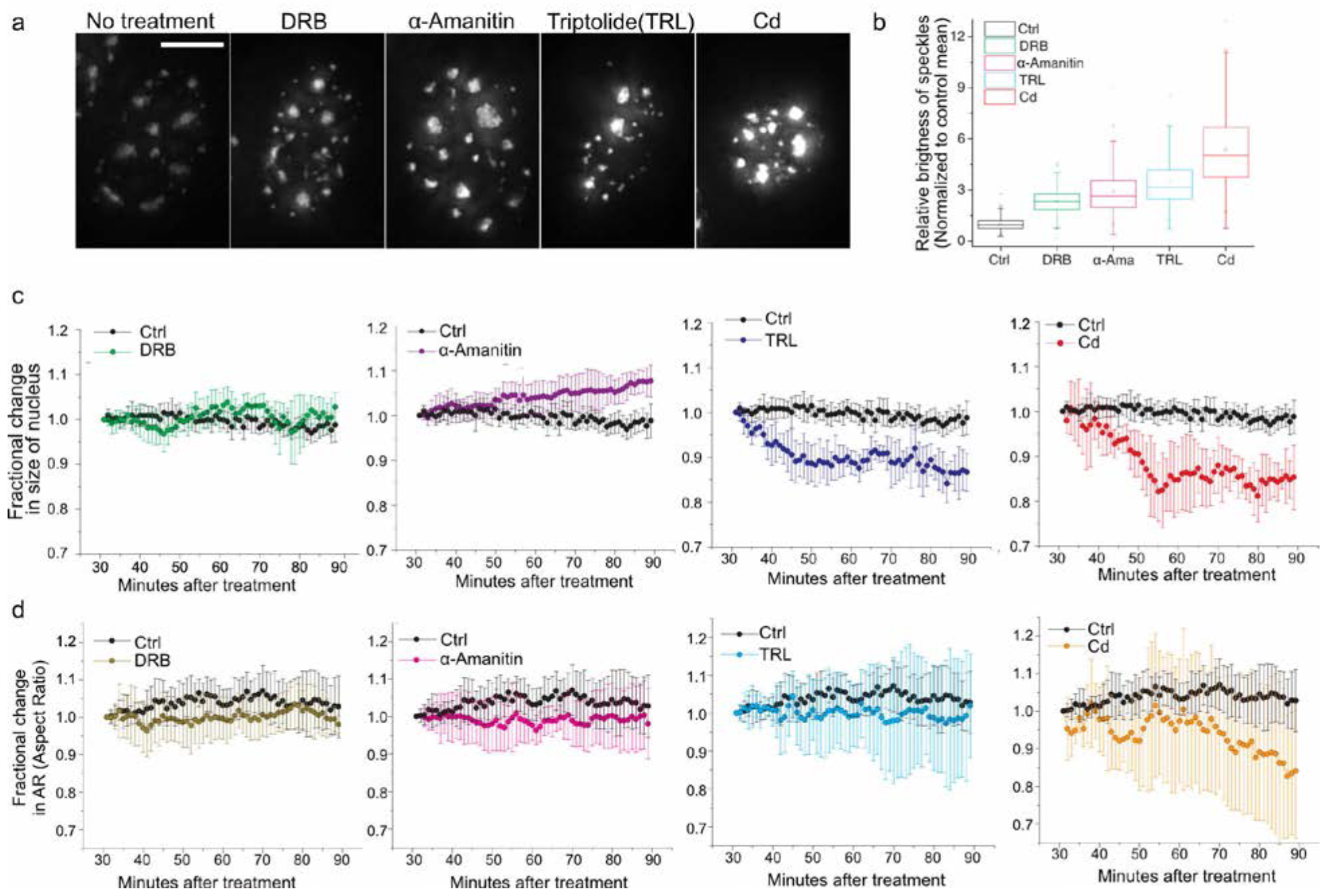
Morphological changes of nuclear speckles and nuclei after treatment with different transcription inhibitors or cadmium (Cd). **a.** Fixed cell images after 2 hr treatment with the indicated chemical. Scale bar: 5um. **b.** Box plots summarizing statistical distribution of relative brightness of individual speckles after treatment with indicated chemical normalized to the mean of speckles in control cells. **c.** Fractional change in the size of nucleus during 1 hr live cell imaging of control (Ctrl, black) or treated cells (Chemical indicated, color). Triptolide (TRL). Error bars represent standard deviations between cell nuclei. **d.** Fractional change in aspect ratio of the nuclear major axis to minor axis during 1 hr live cell imaging. Error bars represent standard deviations between cell nuclei.

**Supplementary Figure. 2.**
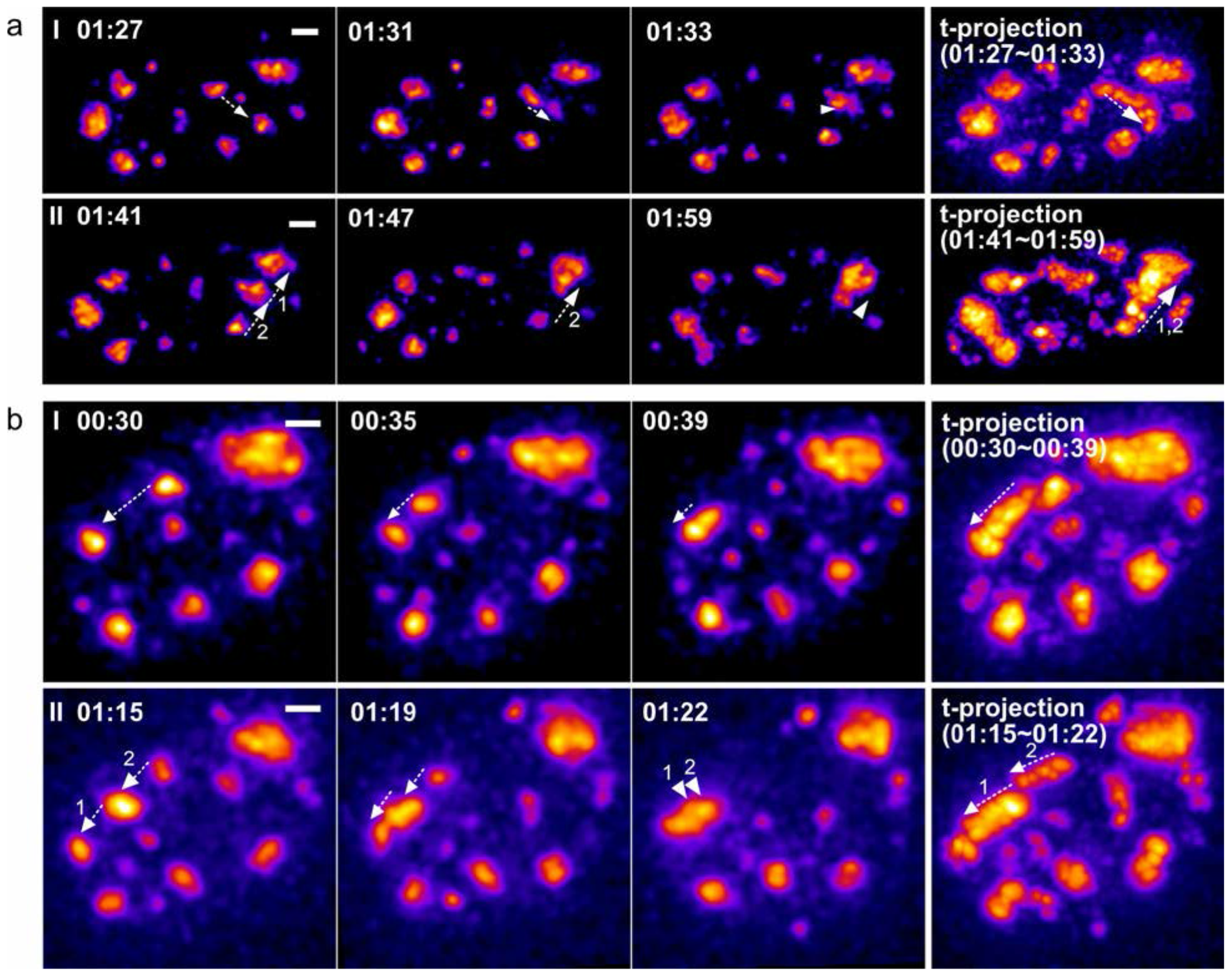
Linear path of long-range (>1μm) directional speckle movements. SON-GFP images represent maximum intensity 2D projections (left) of 3D image stacks or maximum intensity projections over time (t-projection) of these 2D projections (right). Time (hr:min) is after DRB treatment. Scale bars: 1 μm. **a.** (1,11) Examples of long-range directional speckle movements within one nucleus. (Left) White arrows: direction of each speckle movement. **b.**(I,II) Additional examples from second nucleus.

**Supplementary Figure. 3.**
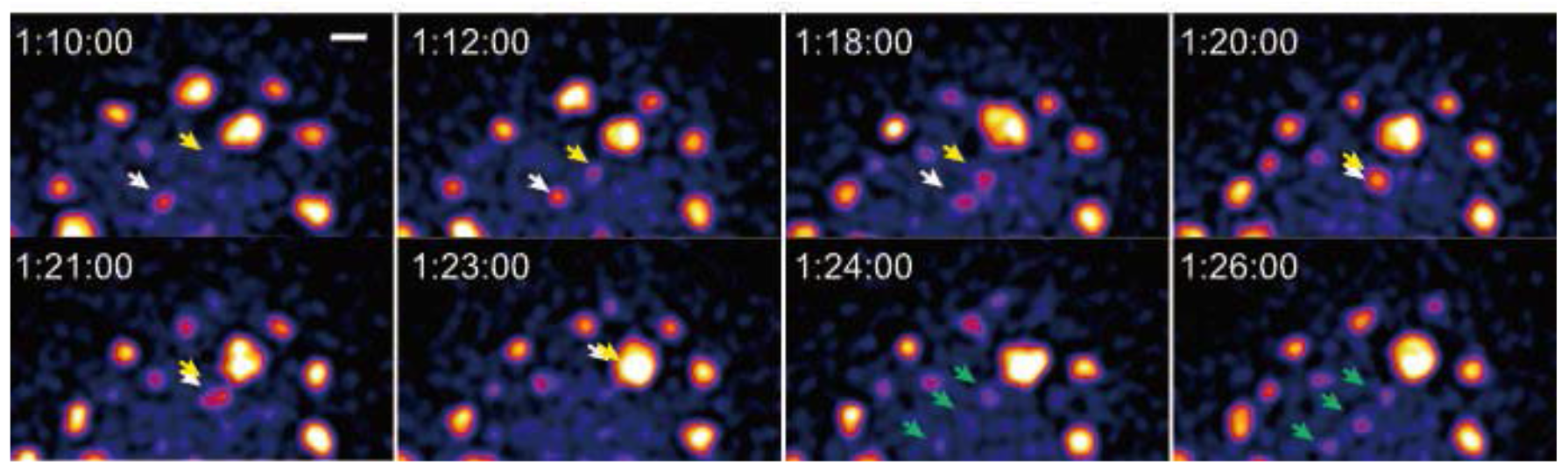
Nucleation of speckles along a path followed by a nuclear speckle. SON-GFP images represent maximum intensity 2D projections of 3D image stacks. Time (hr:min:sec) is after DRB treatment. Scale bar: 1 um. As a first speckle (white arrowhead) moves towards a large, target speckle, a second, small speckle (yellow arrowhead) appears to nucleate at a position on linear path connecting the first and target speckles. The first speckle moves along this linear path, fuses with the second speckle, and then the fused speckle continues along this linear path and fuses with the large target speckle. One minute later, three new small, collinear speckles (green arrowheads) form along this same path.

**Supplementary Figure. 4.**
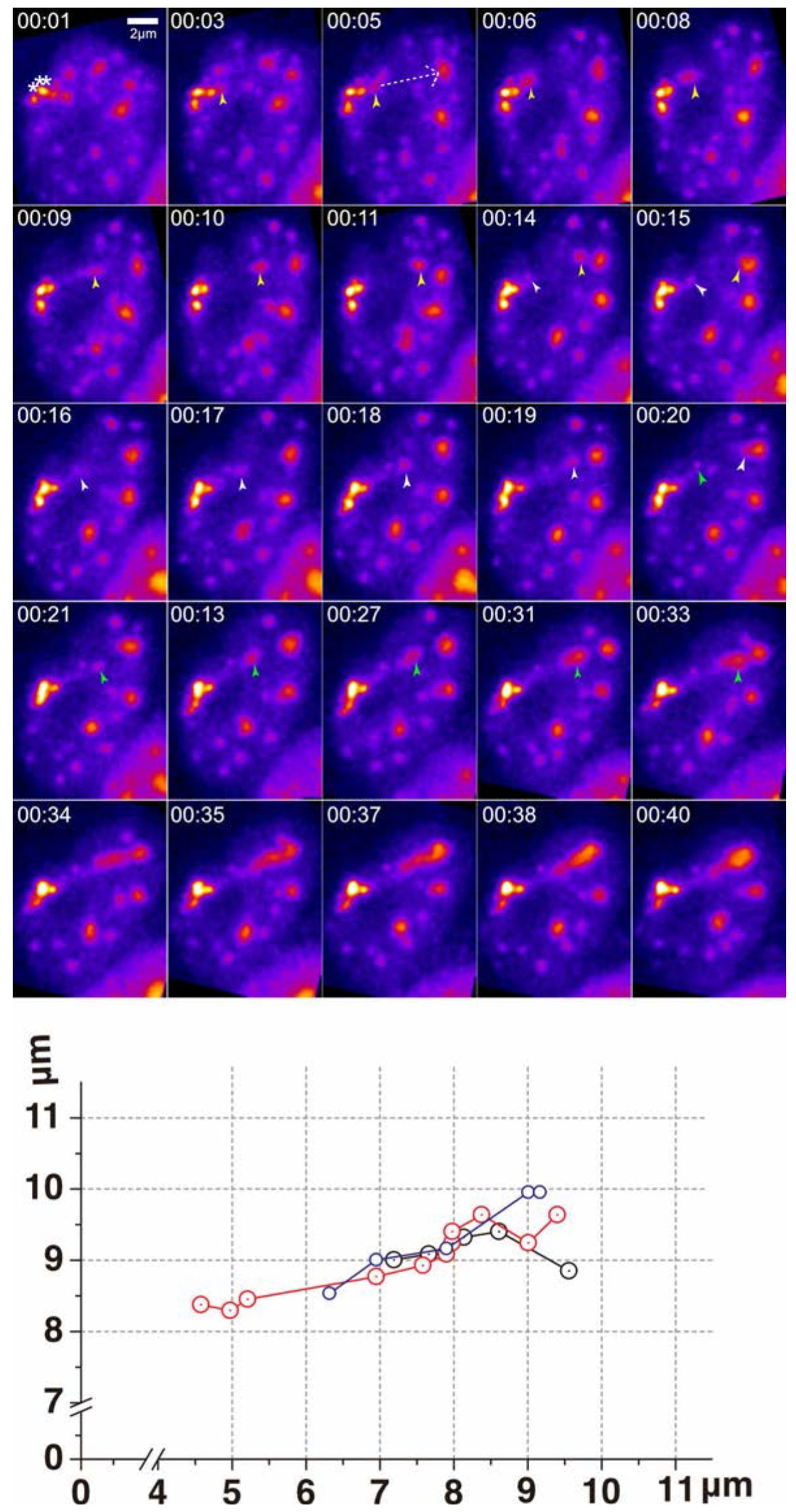
Repeated cycles of speckle formation, translocation and fusion. Top: Pseudo-colored GFP images showing SON-GFP (speckles) and GFP-lac repressor (Hsp70 plasmid transgene array) (Khanna et al. 2014) represent maximum intensity 2D projections of 3D image stacks. Time (hr:min) is after heat-shock induction of Hsp70 transgene array. Scale bar: 2 μm. Nuclear speckles nucleate adjacent to Hsp70 transgene array (brightest spot, marked by asterisks 1min), move along linear path (white dashed arrow, 5 min) to fuse with large target nuclear speckle. Three cycles of speckle nucleation, long-range movement, and fusion to same target speckle are seen. Arrowheads (yellow, white, and green- in order of speckle formation) mark moving speckles. Bottom: Trajectories (x,y) of each long range directional speckle movement.

**Supplementary Figure. 5.**
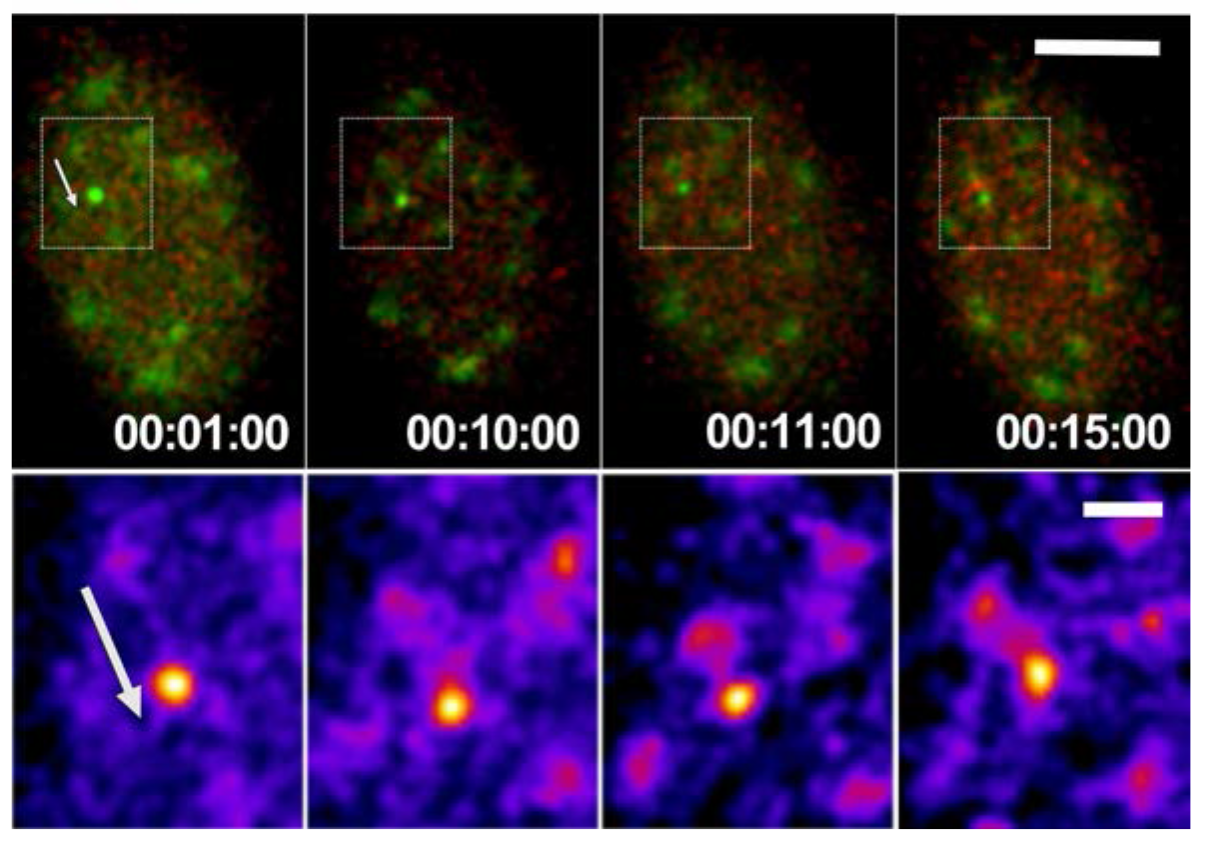
Nuclear speckle movement towards heat-shock, transcriptionally activated Hsp70BAC transgene occurs along path marked by local GFP-SON accumulation. Top: Speckle movement (white arrow) occurs along a linear path. Images of SON-GFP (speckles, light green) and GFP-lac repressor (Hsp70 transgene, brighter green) and mCherry-MS2-binding protein (red) represent 2D maximum intensity projections of 3D image stacks. Time (hr:min:sec) is after heat-shock induction of Hsp70 transgene. The mCherry-MS2-binding protein stains MS2-tagged transcripts of the Hsp70 transgene. Scale bar: 5μm. Bottom: Enlarged views of boxed region from top images. Subsequent speckle movement occurs along linear path (white arrow) marked by GFP-SON accumulation at 1:00 min after heat-shock. Pseudo-color display of GFP intensities provides improved dynamic range for visualization. Images were coded with ‘Fire’ LUT in ImageJ. The brightest spot (yellow) marks the Hsp70 transgene; purple regions are GFP-SON. Scale bar: 1 μm.

